# Parallel Multilink Group Joint ICA: Fusion of 3D Structural and 4D Functional Data Across Multiple Resting fMRI Networks

**DOI:** 10.1101/2024.03.21.586091

**Authors:** K M Ibrahim Khalilullah, Oktay Agcaoglu, Jing Sui, Marlena Duda, Tülay Adali, Vince D Calhoun

## Abstract

Multimodal neuroimaging research plays a pivotal role in understanding the complexities of the human brain and its disorders. Independent component analysis (ICA) has emerged as a widely used and powerful tool for disentangling mixed independent sources, particularly in the analysis of functional magnetic resonance imaging (fMRI) data. This paper extends the use of ICA as a unifying framework for multimodal fusion, introducing a novel approach termed parallel multilink group joint ICA (pmg-jICA). The method allows for the fusion of gray matter maps from structural MRI (sMRI) data to multiple fMRI intrinsic networks, addressing the limitations of previous models. The effectiveness of pmg-jICA is demonstrated through its application to an Alzheimer’s dataset, yielding linked structure-function outputs for 53 brain networks. Our approach leverages the complementary information from various imaging modalities, providing a unique perspective on brain alterations in Alzheimer’s disease. The pmg-jICA identifies several components with significant differences between HC and AD groups including thalamus, caudate, putamen with in the subcortical (SC) domain, insula, parahippocampal gyrus within the cognitive control (CC) domain, and the lingual gyrus within the visual (VS) domain, providing localized insights into the links between AD and specific brain regions. In addition, because we link across multiple brain networks, we can also compute functional network connectivity (FNC) from spatial maps and subject loadings, providing a detailed exploration of the relationships between different brain regions and allowing us to visualize spatial patterns and loading parameters in sMRI along with intrinsic networks and FNC from the fMRI data. In essence, developed approach combines concepts from joint ICA and group ICA to provide a rich set of output characterizing data-driven links between covarying gray matter networks, and a (potentially large number of) resting fMRI networks allowing further study in the context of structure/function links. We demonstrate the utility of the approach by highlighting key structure/function disruptions in Alzheimer’s individuals.

## Introduction

Independent component analysis (ICA) strives to disentangle mixed independent sources. ICA has shown great promise in the analysis of fMRI data and is a widely used tool. When applied to fMRI data, ICA can effectively segregate sources that are independent either spatially or temporally, performing well under appropriate assumptions (Calhoun et al., 2001b; Calhoun & de Lacy, 2017; Calhoun et al., 2021; T. Adali, 2014). ICA is also used for multimodal fusion, allowing joint analysis of multiple modalities that can enhance our comprehension of the human brain and its disorders, offering a more comprehensive perspective compared to studies focusing on a single modality (Calhoun & Sui, 2016; Sendi et al., 2020). Employing multiple modalities can harness the complementary information they offer, enhancing the overall depth of understanding. However, multimodal fusion is also a challenging task, as it requires combining data with different properties, dimensions, and scaling. An additional complexity is that multiple datasets from an individual are likely not statistically independent and identically distributed.

The current landscape of multimodal neuroimaging research has seen notable contributions, yet challenges persist in seamlessly integrating information from multimodal data. (Luo et al., 2020) demonstrated a structural–functional relationship using a dataset comprising over 1500 individuals, employing a two-step process with separate analyses rather than a joint approach. In a similar vein, (Sendi et al., 2020) investigated the transition from a normal brain to very mild Alzheimer’s disease (vmAD) through separate multimodal analyses. The joint analysis of multiple imaging data types from the same individual has been shown to be particularly valuable in this context. (Qi et al., 2022) innovatively developed a three-way parallel group ICA fusion method to distinguish between schizophrenia and control subjects while linking data together with different dimensionality. Additionally, (Duan et al., 2020) introduced a multimodal fusion approach known as aNy-way ICA, capable of detecting interconnected sources across any number of modalities without imposing orthogonality constraints on the sources.

Many multimodal approaches operate under the assumption that various modalities originate from a shared distribution. Additionally, in these approaches, the high-dimensional fMRI data are often condensed into a singular map for each subject, for example, the amplitude of low frequency fluctuation (ALFF) (Hare et al., 2017; Jia et al., 2021; Turner et al., 2012) or a single intrinsic network like default mode. The use of a highly summarized measure like the ALFF map is quite lossy and may not capture the full spatial complexity of the fMRI data. This reduction can lead to a loss of detailed spatial information, making it challenging to integrate with other modalities that might benefit from more nuanced spatial representations. Our previous model called parallel multilink joint ICA (pml-jICA) provides a powerful approach that allows us to fuse gray matter maps to multiple functional networks simultaneously while also allowing for different distributions (Khalilullah et al., 2023). Our pml-jICA approach focused on multiple brain networks but did not fully capture the inter-relationship among brain networks nor did it optimize for individual sets of linked networks and gray matter. This paper aims to bridge the existing gap in integrating multimodal MRI data by extending ICA as a unifying framework for multimodal fusion. We developed a novel approach that combines concepts from Group ICA and parallel multilink joint ICA, to fuse information about multiple brain networks, their temporal information, and links to brain structure via gray matter. As we show, this represents a powerful approach that allows network specific fusion to gray matter, estimation of functional network connectivity (FNC) within the fused model, and network specific gray matter decompositions. The method is called parallel multilink group joint ICA (pmg-jICA). The effectiveness of the group ICA hinges on the separability of the estimated mixing matrix among subjects. The proposed approach allows us to leverage back-reconstruction to estimate spatial maps for individual brain networks from a functional-structural fusion algorithm which can be individually tested for group difference or association with a variable of interest. We apply our approach to an Alzheimer’s dataset resulting in linked structure-function output including a total of 53 brain networks from our NeuroMark spatially constrained ICA pipeline (Du et al., 2020).

### Model Development

We introduce the proposed parallel multilink group jICA model in Figure-1. The analysis stages of this model include: a) preprocessing, b) designing data matrix and data reduction, c) joint independent component estimation, and d) z-scored filtering and interpretation of results. Our proposed model estimates joint spatial map and subject loading from gray matter (GM), denoted as 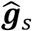; and intrinsic connectivity networks (ICN), denoted as 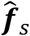; *where, s* = 1, 2 …, *N*; *s* represents subject number. The first stage (a) represents transformation **T**(.) of the raw data, mainly including reslicing and smoothing. The number of ICNs can be arbitrary, but here we include 53 ICNs, indexed as 1, 2, …, *k*; *k* = 53, from our NeuroMark pipeline (Du et al., 2020) in the subsequent application of our new approach. The second stage (b) consists of designing the data matrix and dimensionality reduction. Data from the fMRI modality were organized into 53 distinct matrices, one for each of the 53 ICNs, containing spatial maps from all subjects for that particular ICN. Data reduction for these combinations is performed in two steps, first one on data from each matrices 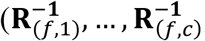, *where C* = 1,2, …, c) for the ICNs of all matrices, *C* represents number of distinct matrices, c is the 53rd ICN, and second one on the concatenated whitening signals from the first steps 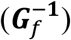. It is used to reduce the computational load of simply taking all subjects’ data and identify common spatial patterns across a group of individual ICN for all subjects. All gray matter (GM) images are reduced in one step, demonstrated in Figure 1. The third stage (c) involves joint independent component estimation. Since, the GM and ICN come from different distributions, we use our previous algorithm parallel multilink joint ICA (pml-jICA) (Khalilullah et al., 2023), which performs alternating initialization and estimation, thus relaxing the original jICA assumption of same distribution for all modalities. This also prevents one modality from dominating the other in the estimation of the maximally independent components. The fourth stage (d) consists of filtering the resulting spatial maps and estimating subject loadings. In the figure, *d* represents number of independent sources. Finally, we reconstructed network-specific maps and loadings using dual regression (Erhardt et al., 2011).

**Fig. 1:**
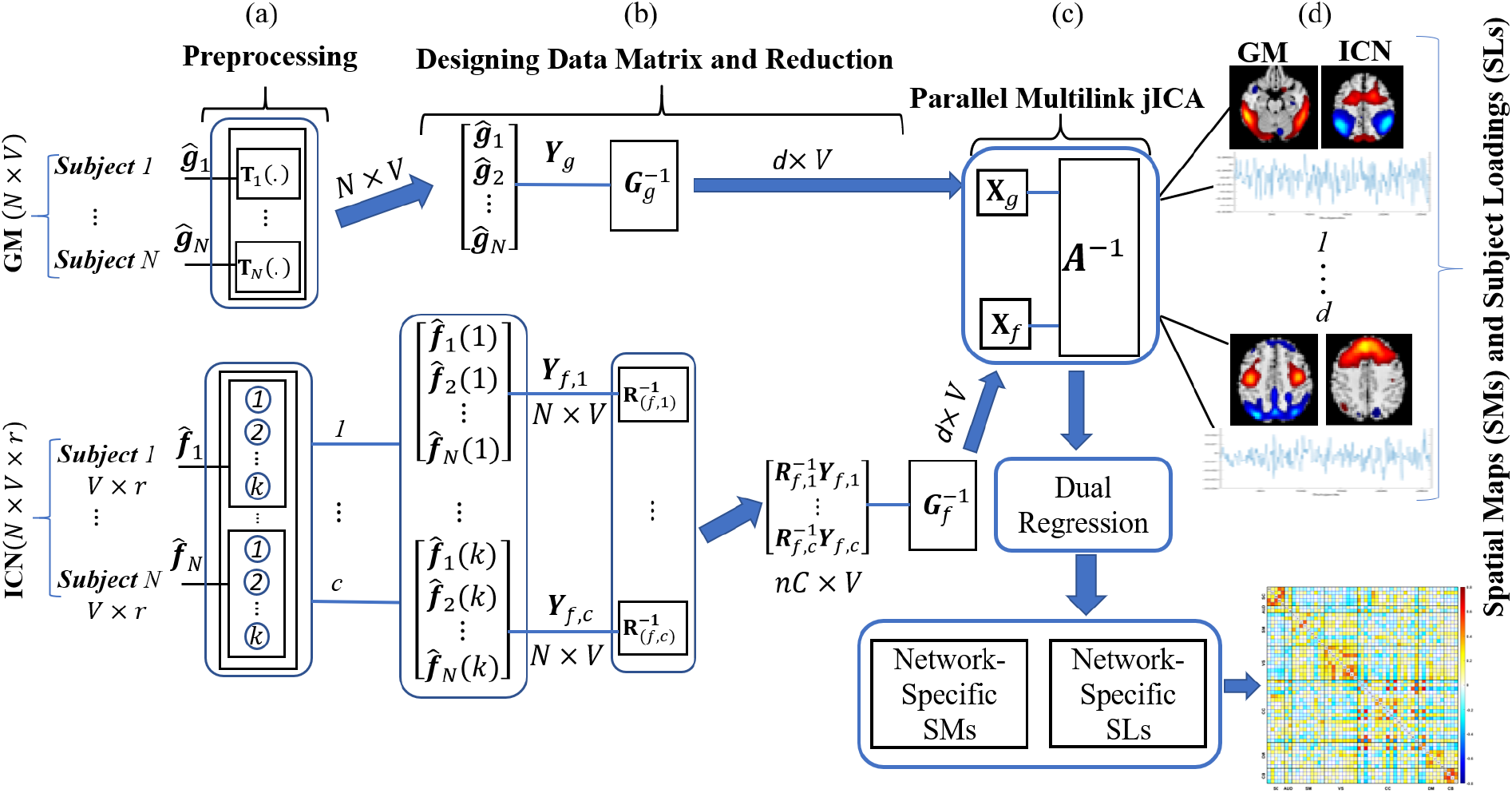
Model for Parallel Multilink Group jICA allows linking of an arbitrary number of resting networks with gray matter maps, and resulting testing of loading parameters, voxel values, and functional network connectivity maps; the states (a)-(d) are explained in the above section.

### Mathematical Formulation

In this model, data from the GM maps and their corresponding ICNs are first subjected to data reduction across all GM maps, then concatenated in the reduced dimension. This concatenated matrix is then further reduced in a second data-reduction stage. The resulting matrix can be used as an input of the ICA estimation stage. The spatial maps for each ICN are then back-reconstructed from the aggregated mixing matrix. In our prior work, we showed that the individual unmixing matrices will be approximately separable across subjects and the back-reconstructed data will be a function of primarily the data within subjects rather than across subjects (Calhoun et al., 2001a), which is the basic foundation of our model.

#### ICA Estimation

Let, 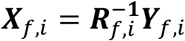 be the *n* × *V* reduced data matrix of ICN from combination *i*, where ***Y***_*f*,*i*_ is the *s* × *V* data matrix, 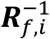 is the *n* × *s* reduction matrix of the ICN, which is computed via a PCA decomposition. The reduced data of the ICN from all combination are then concatenated into a matrix and reduced this matrix to the number of components, *d*, to be estimated. The *d* × *V* reduced matrix (second stage reduction) of all the matrices is

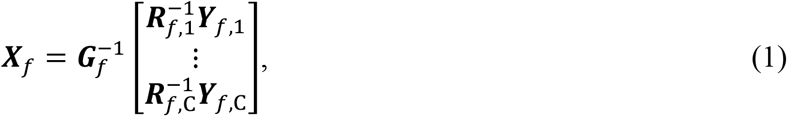

where 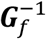 are the two *d* × *nC* dimension reduction matrices for fMRI. All GMs are reduced in one step, let, 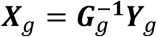 be the *d* × *V* reduced data matrix of all GMs, where ***Y***_*g*_ is the *s* × *V* data matrix after the preprocessing stage, 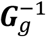 is the *d* × *s* reduction matrix, which is computed via a PCA decomposition.

In the ICA estimation, we can write

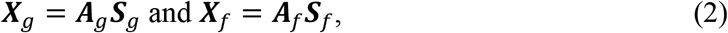

where ***A***_*g*_ and ***A***_*f*_ are *d* × *d* mixing matrices and ***S***_*g*_ and ***S***_*f*_ are the *d* × *V* independent source maps of the GM and ICN, respectively.

The joint mixing matrix, ***A***, is estimated from the ***A***_g_ and ***A***_f_ using parallel multilink joint ICA (pml-jICA) (Khalilullah et al., 2023). Thus, the equation becomes

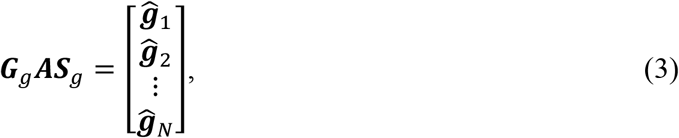

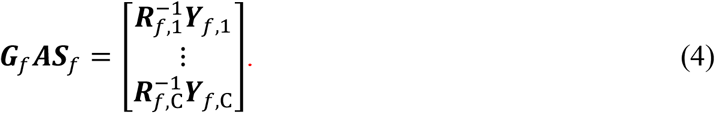

#### Back-reconstruction

Dual regression is an indirect back reconstruction approach using least square to estimate subject-specific loadings and spatial maps. This approach first estimates time courses using aggregated ICA maps as regressor, then subject specific spatial maps were estimated using the timecourses as regressor (Erhardt et al., 2011). In this joint analysis, we first estimate network-specific loadings, then ICN level joint spatial maps using the aggregated joint source maps from parallel multilink group jICA (pmg-jICA). For each combination, *i* = 1, …, *c*, let the input multimodal data be the product of network-specific subject loadings (SLs) and spatial maps (SMs), ***F***_*i*_ and ***S***_*i*_, plus error, ***ε***_*i*_,

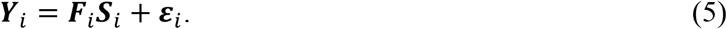

The highly correlated components are grouped into the same component in the ICA analysis. Therefore, variation among networks may have network specific SMs that are a mixture of a few aggregate SMs. In dual regression based back-reconstruction, the first assumption is that all ICNs share a common SM, ***S***_*i*_ ***≡ S***, *i* = 1, …, *C*.

Then, from ICA in equation (2), regression expression becomes,

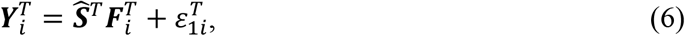

where 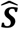 is an aggregate joint SMs from equation (2), 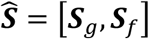, and the estimated joint mixing matrix, ***A***.

Least square estimation for the joint 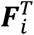 gives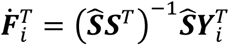, or the transpose (Calhoun et al., 2004),

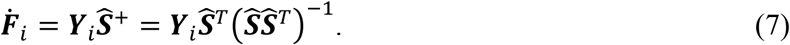

It is noted that the product of the joint aggregate SM and ICN specific loading is a perpendicular projection of the data onto the column space of 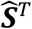, denoted as 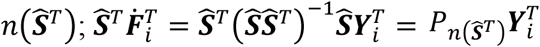.

Thus, the original assumption of common SMs is relaxed for network-specific joint spatial maps estimation. Based on the estimated 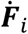, let

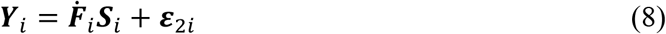

with *E*[***ε***_2*i*_] = 0. Least square estimation for the network-specific spatial maps, ***S***_*i*_, gives

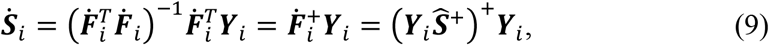

where the ‘+’ sign represents Moore-Penrose pseudoinverse.

The product of each network-specific subject loading (SL) and spatial map (SM) gives

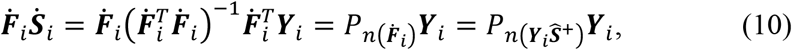

where 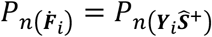 is a perpendicular projection onto the column space of 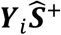, denoted as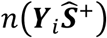.

This paper utilizes experimental data derived from the longitudinal Open Access Series of Imaging Studies (OASIS-3). The data collection spanned 15 years and originated from various ongoing studies conducted at the Washington University Knight Alzheimer Disease Research Center (Pamela J. LaMontagne, 2019). The dataset comprises MRI and PET imaging, along with relevant clinical data for a total of 1098 participants. Our analysis focused on utilizing sMRI and resting state BOLD sequences. To assess neural activity during rest, participants were instructed to remain quietly with open eyes while two 6-minute resting state BOLD sequences were recorded. The OASIS-3 dataset includes a total of 2165 MR sessions (236 from 1.5T scanner and 1929 from 3.0T scanner) for sMRI with T1w scan type, 1691 MR sessions (2 from 1.5T scanner and 1689 from 3.0T scanner) with BOLD-resting state scan type, and various other scan types. Segmentation of T1w images was performed using statistical parameter mapping (SPM12, http://www.fil.ion.ucl.ac.uk/spm/). For each participant, we employed imaging data, demographics information, and the clinical dementia rating (CDR) scale at any cognitive functionality stage. As per the CDR scale, participants were required to have CDR ≤ 1 during the clinical core assessment. Participants who reached a CDR of 2 were no longer eligible for the study. Among the 1098 participants, 850 initially presented as cognitively normal adults. Out of these, 605 remained normal, while 245 transitioned to cognitive impairment at various stages, with ages ranging from 42 to 95 years. The CDR scale for the remaining 248 participants exceeded zero.

The fMRI data underwent preprocessing using SPM12. Initially, rigid body motion correction followed by slice-timing correction was conducted to address subject head motion and timing disparities in slice acquisition. Subsequently, the fMRI data were normalized to the Montreal Neurological Institute (MNI) standards and resliced into 3 mm × 3 mm × 3 mm isotropic voxels. These resliced images were further subjected to smoothing using a Gaussian kernel with a full width at half maximum (FWHM) of 6 mm. The analysis phase involved the identification of 53 intrinsic connectivity networks (ICNs) using the NeuroMark pipeline—a fully automated, spatially constrained independent component analysis (ICA) approach (Du et al., 2020). Spatial priors were established using the Neuromark_fMRI_1.0 component templates. Simultaneously, sMRI data underwent segmentation and spatial normalization using the SPM unified segmentation approach, followed by Gaussian smoothing with a FWHM of 6 mm. For quality control, gray matter (GM) images were randomly grouped, and scans were flagged if the spatial correlation with the group average images fell below 0.95. This quality control process refined the dataset, reducing the initial 1098 subjects to 784, comprising 648 healthy controls (HCs) and 136 individuals with Alzheimer’s disease (AD). We conducted an analysis on a conclusive subset, randomly choosing 260 participants from both the healthy control (HC) and Alzheimer’s disease (AD) groups within the refined dataset. This selection ensured equal sample sizes for both the control and patient cohorts. HCs were defined as cognitively normal with a Clinical Dementia Rating (CDR) score of 0, while AD subjects were categorized as having AD dementia with a CDR ≥ 0.5. The final dataset underwent additional preprocessing, including reslicing and Gaussian smoothing to harmonize voxel sizes, though this step was not strictly required. To facilitate joint analysis, both types of data—sMRI GM and fMRI ICNs—were normalized to unit variance across all subjects within each modality. The demographic and clinical details of the refined experimental data are succinctly presented in Table I.

**Table I:** Demographics and clinical information

| Subset | Metric | Count/Range |
| --- | --- | --- |
| **Full Data** | Total N (HC/AD) | 260 (130/130) |
| **HCs** | Gender (M/F) | 54/76 |
|  | Age at entry (years) | 42.45-94.93 |
|  | CDR Score (range) | 0-0 |
|  | Sum of Boxes (SOB) | 0.0-0.5 |
| **ADs** | Gender (M/F) | 68/62 |
|  | Age at entry (years) | 50.29-95.57 |
|  | CDR Score (range) | 0.5-2.0 |
|  | SOB | 0.5-12.0 |

#### Joint components and loading parameter estimation

The GM and resting fMRI data are preprocessed separately prior to data matrix creation. Next, we generate 53 combinations from each of 130 HC and 130 AD subjects. In the first stage of fMRI reduction, each combination is reduced in dimensionality across subject using principal component analysis (PCA). The concatenated PCA signals of rest fMRI networks are subsequently reduced at the group level, followed by parallel optimization of the weights of the rest fMRI networks and GM to reach convergence using our previous pml-jICA algorithm (Khalilullah et al., 2023), where we also showed that separately learning the two distributions provided performance improvements and avoided one modality dominating the other.

We estimated 10 independent joint components in the final step. Based on the empirical results using an information theoretic approach (Li et al., 2007), the estimated component number was selected and found to separate noise and signals into different independent source maps.

## Results

Results from the reconstructed loadings showed highly significant group differences between AD and HC, also exposed several interesting patterns from the reconstructed source maps that provide network specific coupling between GM and fMRI. In the following section, we present statistically significant network specific loadings after reconstruction with their corresponding independent source maps.

In the following, we present two main sets of results, the first set of results represents eleven GM/fMRI component pairs (out of 10 joint components per ICN/GM combination, or 530 pairs) showing significant group differences. Each combination represents a set of joint GM/ICN components derived from an ICN-GM pairing across all subjects estimated in each group ICA block. The second set of results comes from grouping the output into 10 pairs of 53 ICNs w/ their aggregate GM map, and computing the FNC matrix by cross correlating the loading parameters for the 53 ICNs. This provides insight into the interrelationship among ICNs for a given ICN/GM pairing. The significant joint source maps and FNC computation are demonstrated in Figure 2.

**Fig. 2:**
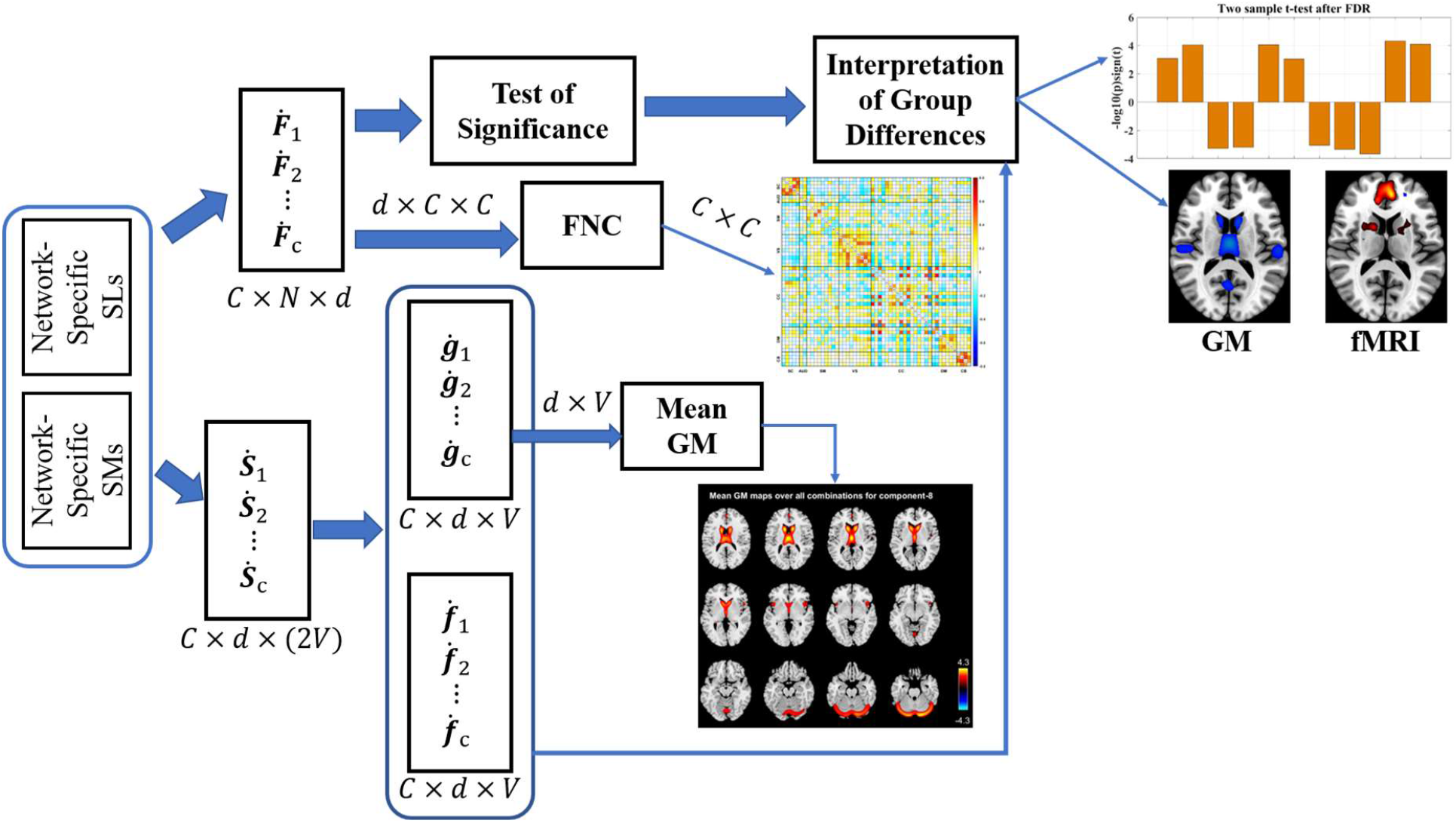
Summary of output from pmg-jICA model including group differences, FNC and Mean GM computation from reconstructed subject maps and subject loading parameters.

### Network Specific Reconstructed Maps

We computed joint reconstructed maps from the aggregated independent joint components for the ICNs and GMs to analyze network specific coupling between GM and fMRI and also to identify differences between HC and AD. The collection of 53 reconstructed joint maps is referred to as ‘53 combinations,’ and in connection with the 53 individual ICN combinations, denoted as com ™ 1, com ™ 2, …, com ™ 53. We computed two-sample t-tests between healthy controls versus patients for each combination. The false discovery rate (FDR) approach was used to correct for multiple comparisons (*q* < 0.05). There are several ICN level significant component based on the threshold for statistically significant, which was set at corrected < 0.001. Figure 3 shows the t-test results of the ICN level significant components and their corresponding logarithmic p-values by group.

**Fig. 3:**
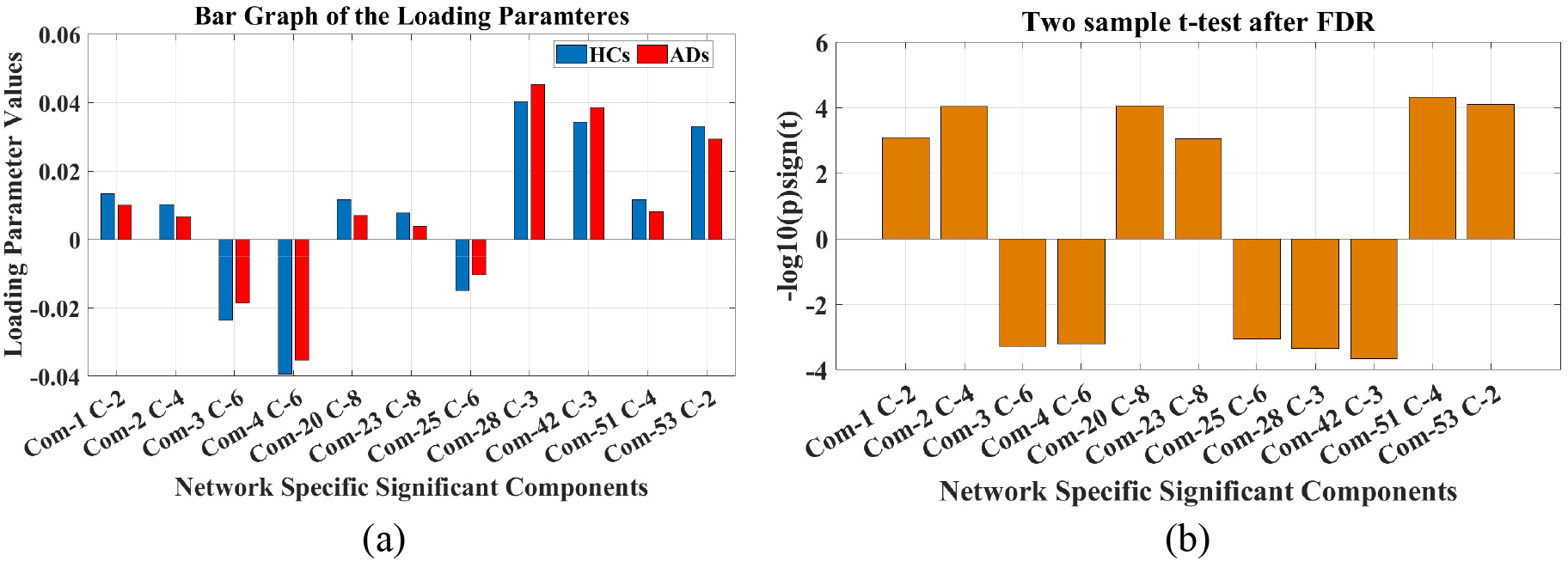
Network specific average loading and statistically significant components; (a) average loading parameter values of the significant components by group; (b) Two-sample t-test results after correction (corrected at *p* < 0.001).

Figures 4-8 demonstrated spatial maps of the significant ICN. Figure 3 indicates that nine components of combinations (1, 2, 3, 4, 20, 23, 25, 51, and 53) contained areas where the absolute value of the joint source values showed reductions in AD and two components of combinations (28 and 42) showed increase in AD, relative to controls. The p-values for the significantly group differing spatial maps and their directions are shown in Figures 4-8. In component 2 of combination 1(Figures 4a-b), the most activated voxels of the GM are mostly in the middle temporal gyrus of the auditory regions, which is increased in AD; most regions in the fMRI are caudate of the subcortical domain and cognitive control areas of the brain, where the activated subcortical regions are lower in AD, but the activated regions of the cognitive control are combination of increases and decreases in AD. In GM of the component 4 of combination 2(Figures 4c-d), insula, inferior frontal gyrus of cognitive control and superior temporal gyrus of AU are lower in AD; in the fMRI thalamus of the subcortical and lingual gyrus of the visual are lower in AD, but some parts of the cognitive control nearly at hippocampal are functionally increased in AD.

**Fig. 4:**
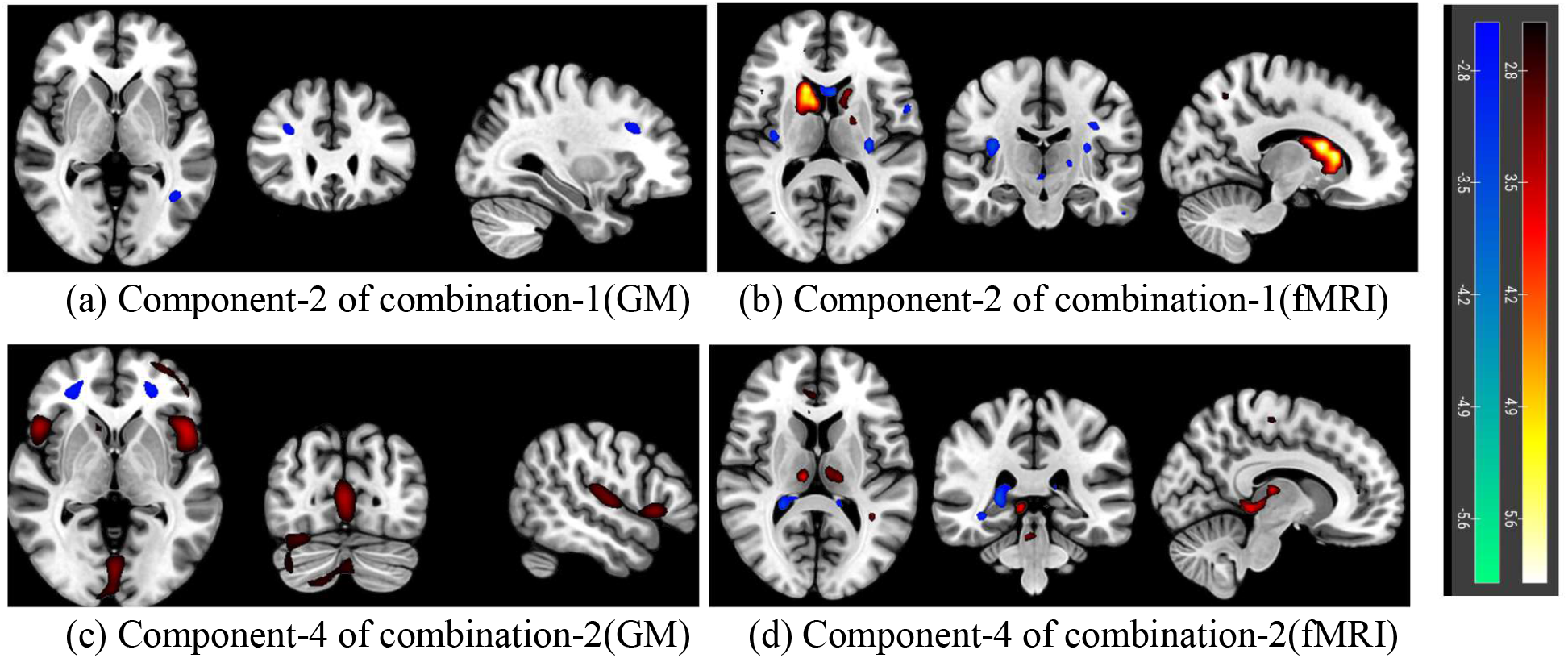
ICN level significant source maps; (a) activated region in GM: middle temporal gyrus, (b) activated region in fMRI: caudate and areas of CC; *p* − *value* = 0.0008 and *HC* > *AD*; (c) activated region in GM: superior temporal gyrus, insula, inferior frontal gyrus, (d) activated regions in fMRI: thalamus and lingual gyrus; *p* − *value* = 0.00009 and *HC* > *AD*.

In component 6 of combination 3 (Figures 5a-b), the significant voxels of the GM are mostly in the thalamus, caudate of the subcortical regions, in addition to auditory (superior temporal gyrus) and cognitive control (insula, inferior frontal gyrus) regions, which are lower in AD. Most activated voxels in the fMRI are located in the superior temporal and middle temporal gyrus of the auditory domain, and subcortical domain and are additive in AD, whereas caudate and putamen as well as parahippocampal regions of the cognitive control domain are lower in AD. One of the strengths of our approach is it can separate overlapping regions into different components, where one is subtractive and another one is additive, functionally and/or structurally. Component 6 of combination 4 (Figures 5c-d) indicates joint relationships between cognitive control GM volume and subcortical and cognitive control fMRI connectivity.

**Fig. 5:**
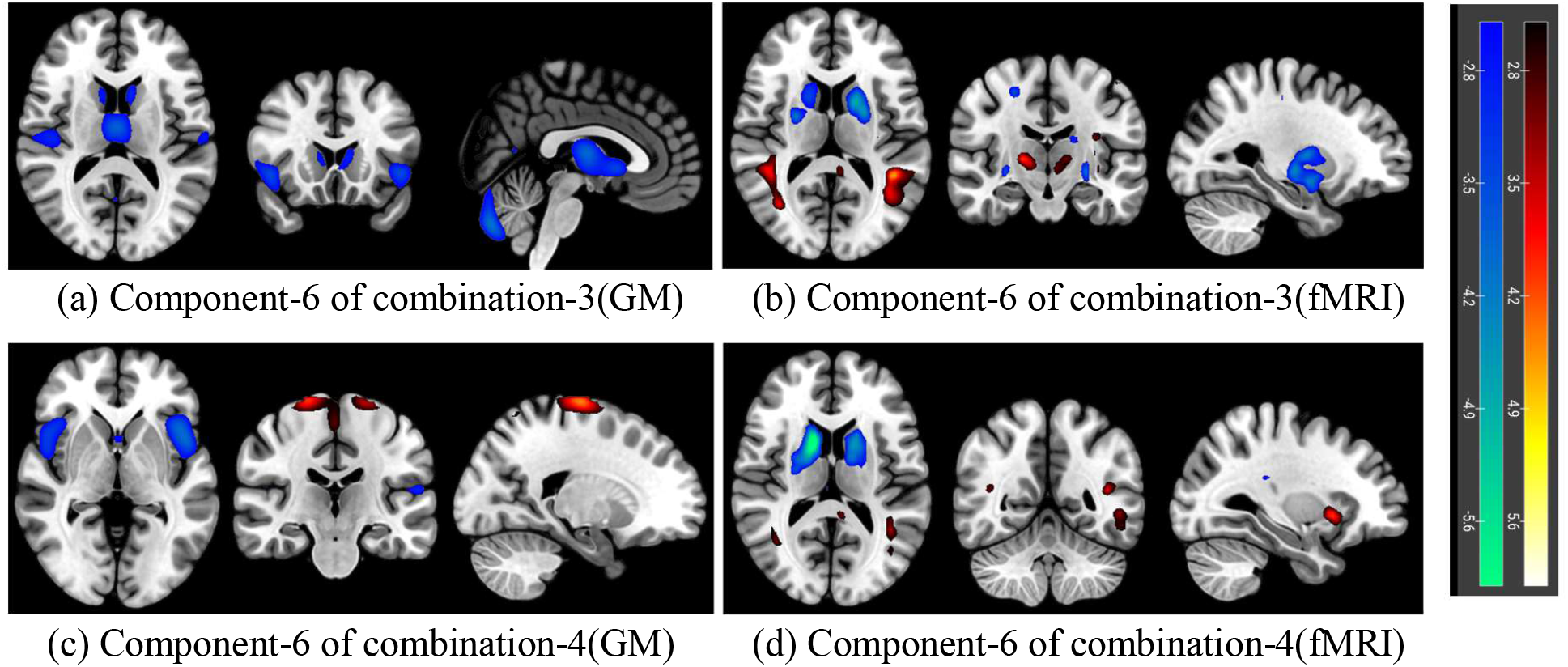
ICN level significant source maps; (a) activated region in GM: superior temporal gyrus, thalamus, caudate, insula, inferior frontal gyrus, (b) activated region in fMRI: superior temporal gyrus, middle temporal gyrus, caudate, putamen, parahippocampal; *p* − *value* = 0.0005 and *HC* < *AD*; (c) activated region in GM: areas of CC, (d) activated regions in fMRI: caudate, putamen; *p* − *value* = 0.0006 and *HC* < *AD*.

Component 8 of combination 20 (Figures 6a-b) highlights joint relationships between subcortical (e.g., caudate, thalamus), cerebellum in GM and visual regions in fMRI, where subcortical (especially caudate and thalamus) and cerebellum regions of the GM are lower in AD, whereas primary visual cortex in fMRI is mostly additive in AD. In component 8 of combination 23 (Figures 6c-d), cerebellum regions in the GM are lower in AD; Visual areas including middle occipital gyrus and auditory areas including middle temporal gyrus are functionally subtractive in AD.

**Fig. 6:**
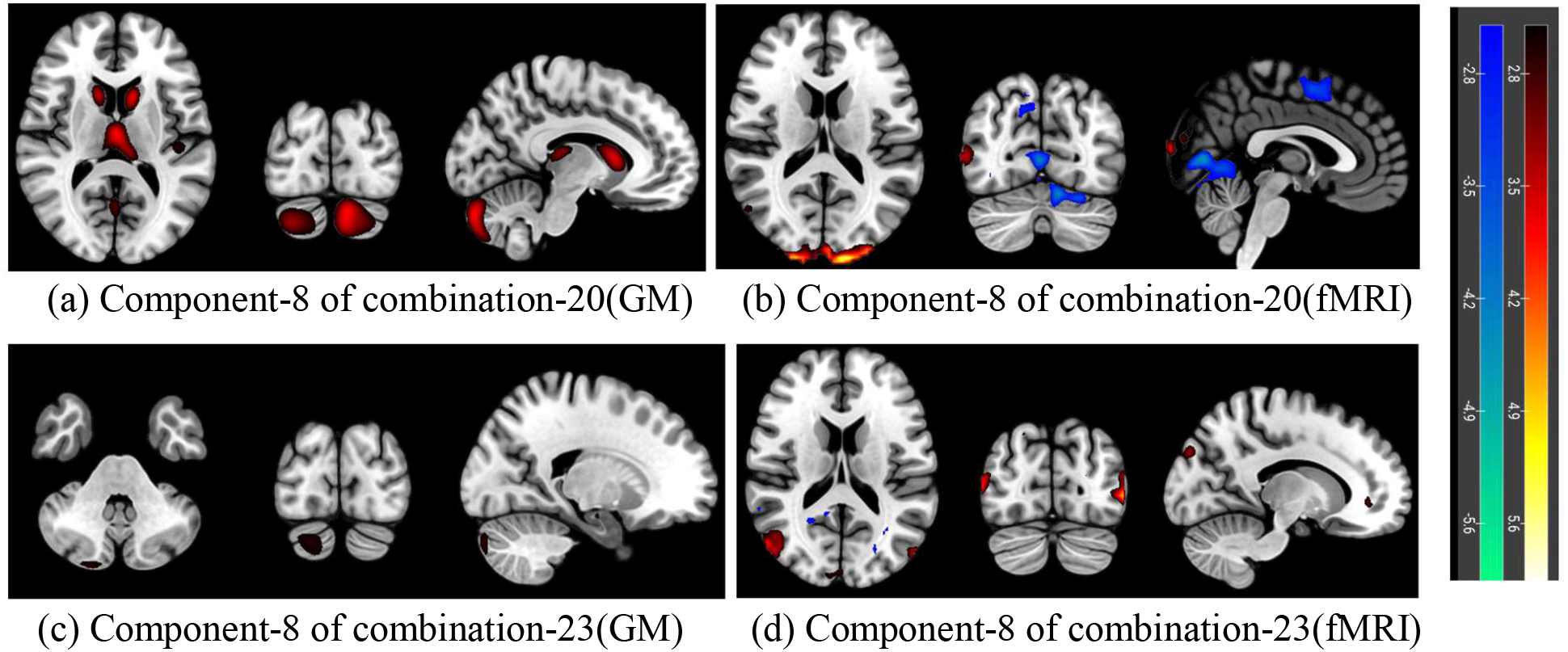
ICN level significant source maps; (a) activated region in GM: caudate, thalamus, areas of CB, (b) activated region in fMRI: areas of VS; *p* − *value* = 0.00009 and *HC* > *AD*; (c) activated region in GM: areas of CB, (d) activated regions in fMRI: middle temporal gyrus, middle occipital gyrus; *p* − *value* = 0.0009 and *HC* > *AD*.

In component 6 of combination 25 (Figures 7a-b), activated voxels of the GM are mostly in the superior temporal gyrus within the auditory domain as well as insula inferior frontal gyrus, and parahippocampal gyrus within the cognitive control domain, all of which are lower in AD; in fMRI, middle temporal gyrus and inferior temporal gyrus in the auditory domain are increased in AD, and some areas of the visual domain are decreased in AD. In component 4 of combination 51(Figures 7c-d), GM cerebellar regions are decreased in AD; whereas in fMRI, we see a combination of increases and decreases, and in addition, the cerebellum is divided into different parts, where one is subtractive and another one is additive; some visual regions in fMRI are increased in AD. In component 2 of combination 53 (Figures 7e-f), activated voxels in the GM are cerebellum areas and inferior temporal gyrus region of the auditory domain, which are lower in AD; most regions in fMRI are cerebellum and visual, where cerebellum is a combination of increases and decreases in AD and visual are increased in AD.

**Fig. 7:**
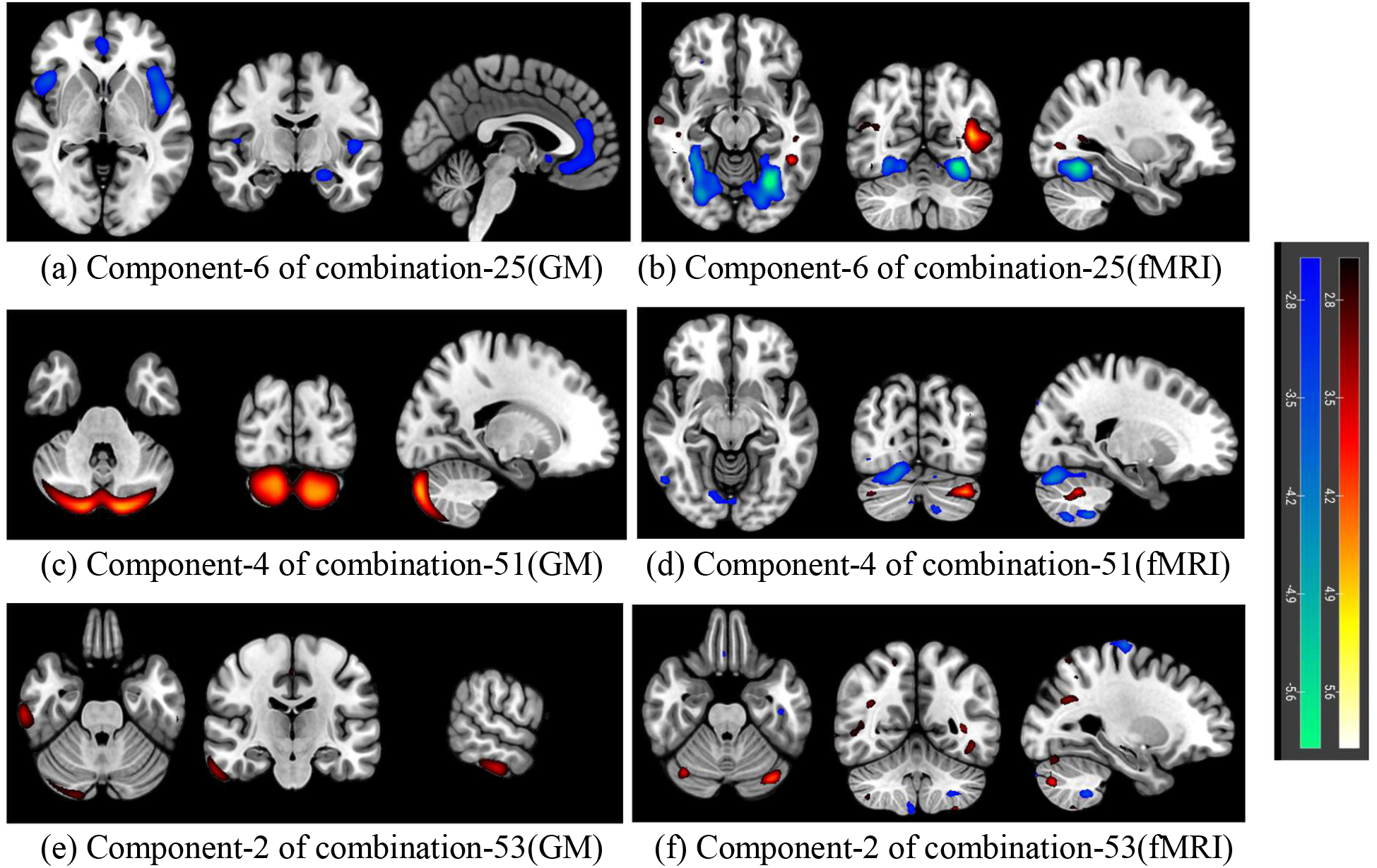
ICN level significant source maps; (a) activated regions in GM: superior temporal gyrus, insula, inferior temporal gyrus, parahippocampal gyrus, (b) activated regions in fMRI: middle temporal gyrus, inferior temporal gyrus, areas of VS; *p* − *value* = 0.0008 and *HC* < *AD*; (c) activated regions in GM: areas of CB, (d) activated regions in fMRI: areas of VS and CB; *p* − *value* = 0.00004 and *HC* > *AD*; (e) activated regions in GM: inferior temporal gyrus, areas of CB, (f) activated regions in fMRI: Areas of VS and CB; *p* − *value* = 0.00007 and *HC* > *AD*.

In component 3 of combination 28 (Figures 8a-b), GM regions include thalamus and caudate within the subcortical domain, superior temporal gyrus in the auditory domain, and some areas of the cerebellum including declive, uvula, and tuber, all lower in AD; in fMRI, superior frontal gyrus and medial frontal gyrus are increased in AD, but middle frontal gyrus are decreased in AD. In component 3 of combination 42 (Figures 8c-d), cerebellum areas of the GM, e.g., cerebellar tonsil, culmin, are increased in AD; in the fMRI, middle temporal gyrus of the auditory and some areas, most likely posterior cingulate, of the default mode network are increased in AD whereas middle frontal gyrus and medial frontal gyrus of the cognitive control and supramarginal gyrus are decreased in AD.

**Fig. 8:**
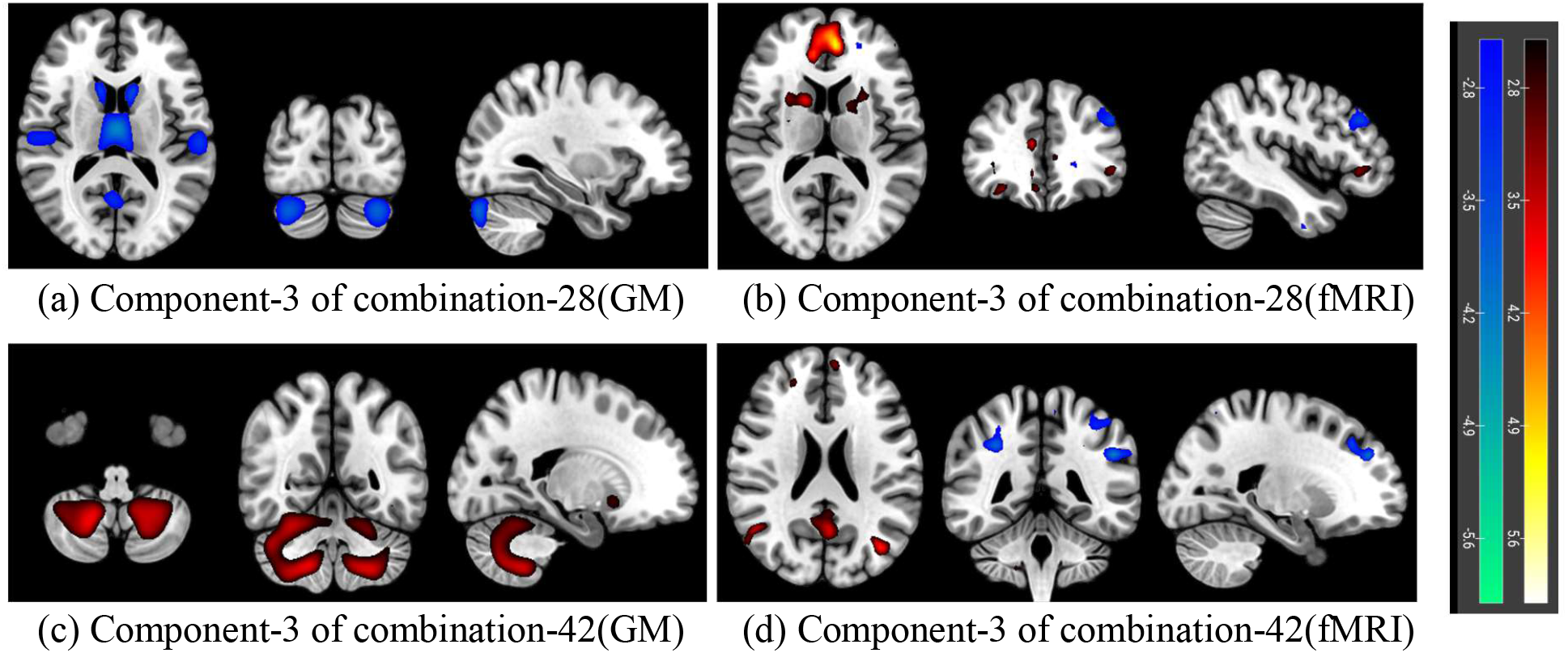
ICN level significant source maps; (a) activated region in GM: superior temporal gyrus, caudate, thalamus, declive, uvula, tuber, (b) activated region in fMRI: superior frontal gyrus, medial frontal gyrus, middle frontal gyrus; *p* − *value* = 0.0004 and *HC* < *AD*; (c) activated region in GM: areas of CB, (d) activated regions in fMRI: middle temporal gyrus, posterior cingulate gyrus, middle frontal gyrus, medial frontal gyrus, supramarginal gyrus; *p* − *value* = 0.0002 and *HC* < *AD*.

The joint loadings provide us with information about group differences between the control and AD groups, as well as spatial maps indicating the implicated areas, which are summarized in Table II and illustrate the structure-function coupling. Additional details are presented in the supplementary materials (Supplementary materials_1). The next section demonstrated functional network connectivity (FNC) that enables is to visualize the coupling between brain networks.

**Table II:**
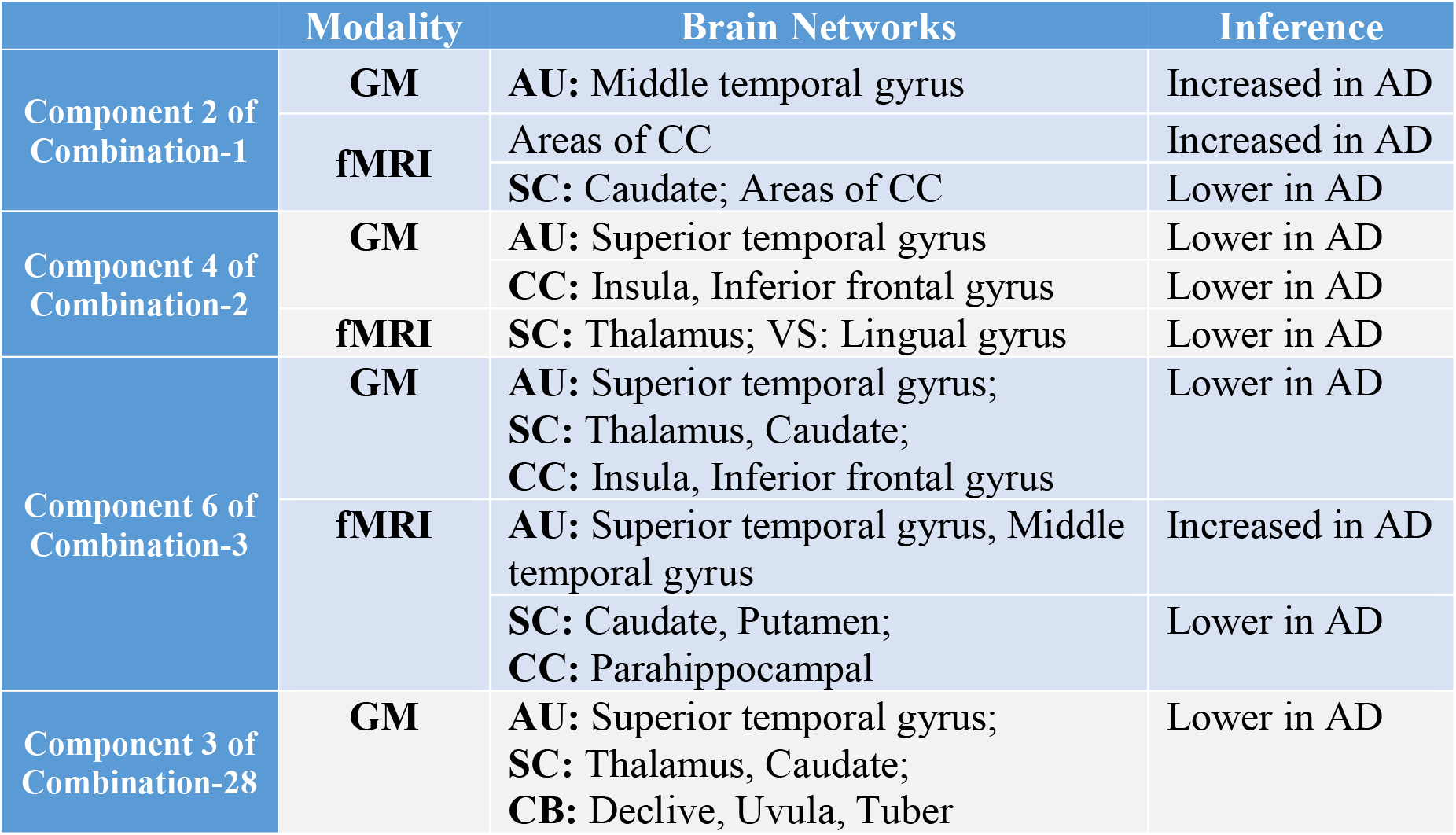
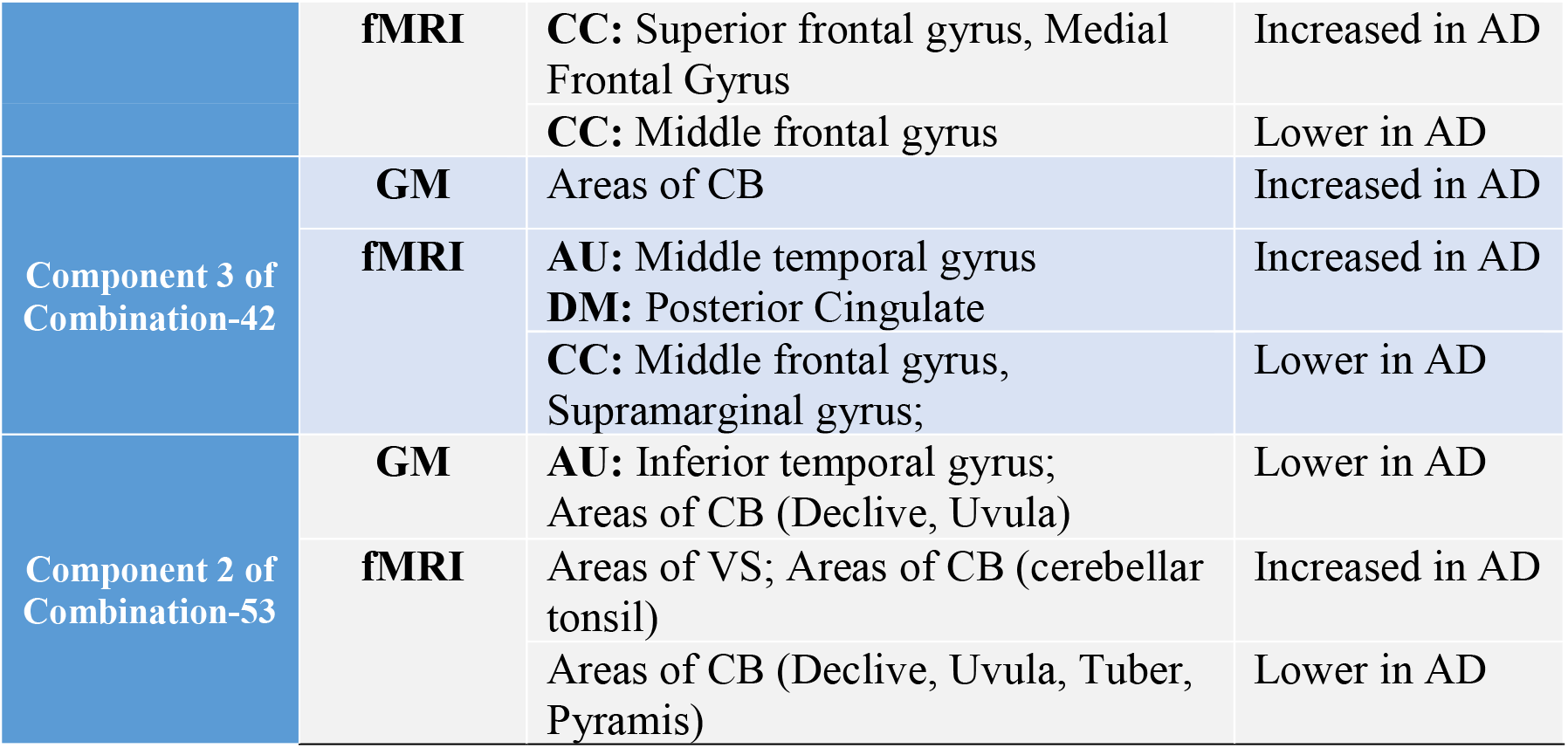
Summarized structure-function correlation

#### Functional Network Connectivity (FNC)

Each reconstructed joint map has a subject loading with the size of *N* × *d*, where *N* = 260 and *d* = 10 represent number of subject and independent components, respectively. From the 53 reconstructed joint loadings, we have generated ten component-wise loadings for all ICNs, which are the 260×53 matrices. Next, we calculated the pairwise linear correlation coefficient between each pair of brain networks, resulting in ten FNC matrices. To streamline presentation, we have highlighted key findings from these matrices, with additional details provided in the supplementary materials (Supplementary Materials_2). These figures illustrate the statistical relationship between two networks. Note that the connectivity strengths show increases or decreases between networks. The ten aggregated GM maps are calculated from the reconstructed gray matter spatial maps by taking average across all combinations (Figure 2).

In Figure 9a, FNC and connectogram of the left image demonstrates some increased functional connections within DM, CB, SC, CC, along with certain components of SM and visual networks. In inter-network connectivity, we observed heightened functional connectivity, denoted by a positive correlation, between specific brain regions. Notably, we detected increased connectivity between the hippocampus of the CC and the postcentral gyrus within the SM network. Furthermore, we identified enhanced connectivity between the precuneus of the DM network and the inferior frontal gyrus of the CC. Additionally, our findings revealed elevated connectivity between the inferior occipital gyrus within the visual network and the inferior parietal lobule of the CC. GM exhibits positive pattern in cerebellum and visual regions of the brain. These regions activate together and share functional and anatomical characteristics that lead to synchronized activity during rest. In the Figure 9c, subcortical, cerebellar and visual regions exhibit the most prominent functional network connectivity within the networks, while the cognitive control domain displays a mixture of connections within intra and inter-networks. In inter-domain connectivity, we found increased FNC between component inferior parietal lobule and superior medial gyrus of CC and precuneus of DM. Furthermore, we identified positive connection between inferior parietal lobule from CC and middle temporal gyrus within AU. The subcortical, cerebellum regions of GM are also structurally connected. Although there isn’t much cross-domain connectivity observed in these figures, the convergence of positive connectivity in both FNC and GM spatial maps for the SC, visual, and cerebellum suggests a strong relationship between the functional and structural aspects of these brain regions. Furthermore, the activation of SC, visual, and cerebellar regions in both GM and fMRI data implies a significant role of the basal ganglia (BG) in facilitating connections across multiple cognitive domains.

**Fig. 9:**
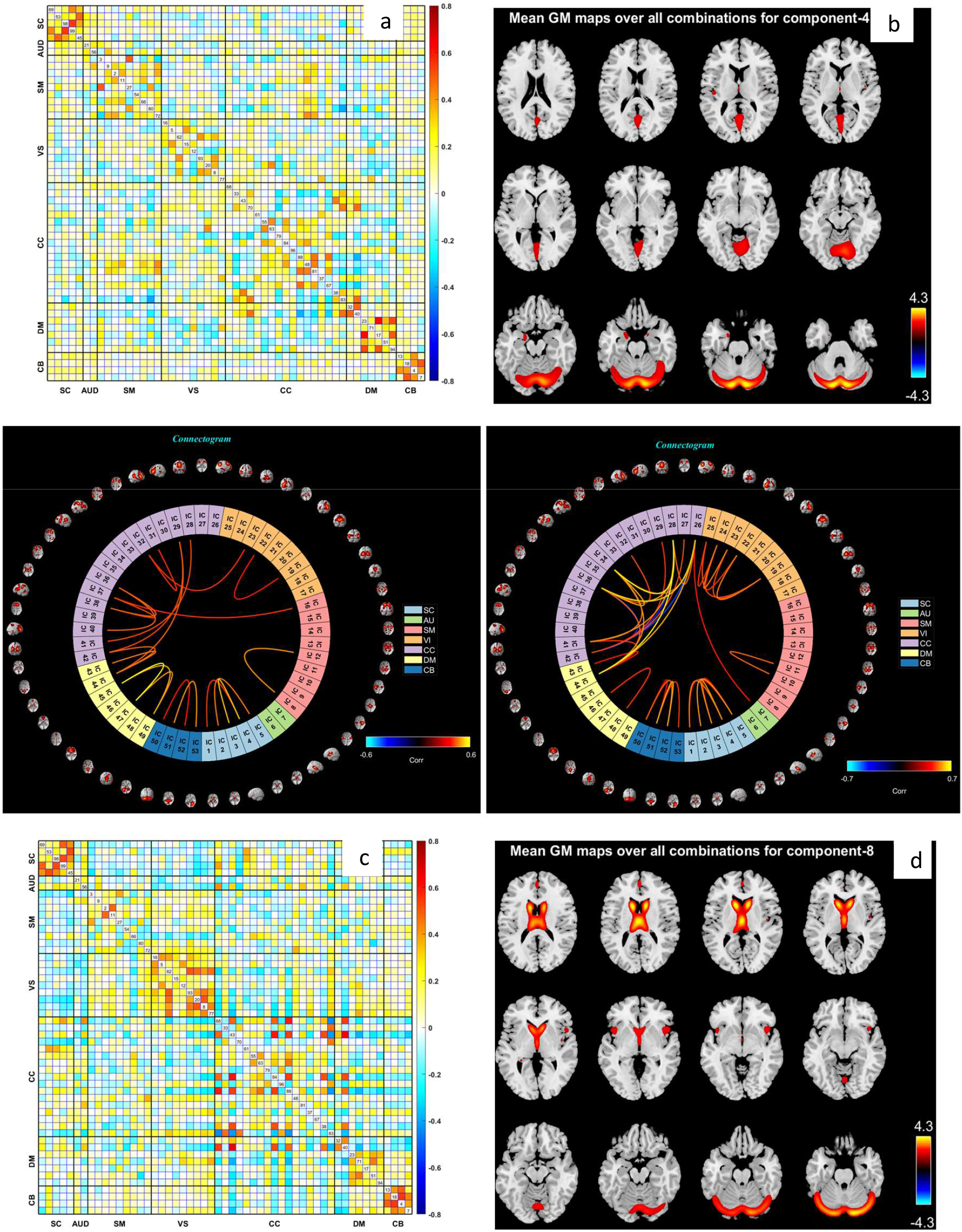
FNC from reconstructed component 4 and 8 of all ICNs; highly correlated regions are (a) hippocampus, inferior parietal lobule, inferior frontal gyrus, middle frontal gyrus, precuneus, postcentral gyrus, paracentral lobule, precentral gyrus, and superior parietal lobule; (c) precuneus, middle temporal gyrus, inferior parietal lobule, superior medial frontal gyrus, supplementary motor area, middle frontal gyrus, inferior frontal gyrus and SC domain; (b) and (d) represent mean GM maps for component 7 and 8, respectively. Connectogram shown in the middle for both FNC, left image for Figure 9a and right image for Figure 9c.

In the context of intra-domain connectivity, FNC exhibits positive connectivity in CB, SC, and VS (Figure 10a). There isn’t extensive cross-connection, except for that observed between the precuneus of the DM and the inferior frontal gyrus originating from the CC. GM also shows positive connectivity in CB, VS, and DM (Figure 10a). Figure 10a and connectogram of the left image suggests that some of networks within SC, CB, and SM are more activated than others.

**Fig. 10:**
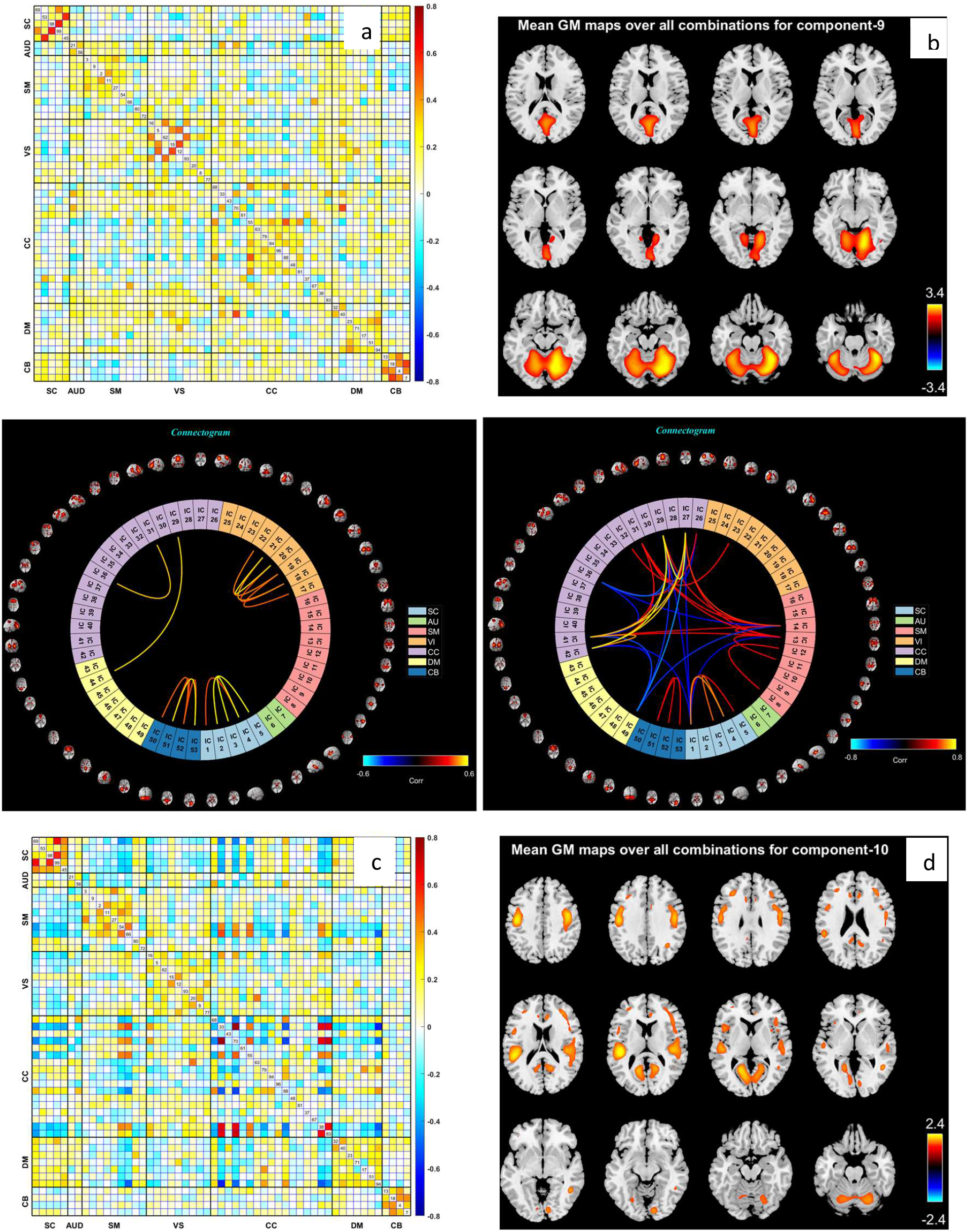
FNC from reconstructed component 9 and 10 of all ICNs; highly correlated regions are (a) hypothalamus, thalamus, superior temporal gyrus, middle occipital gyrus, middle cingulate cortex, insula, inferior frontal gyrus, middle frontal gyrus, posterior cingulate cortex, precuneus, and precentral gyrus, (c) anterior cingulate cortex, posterior cingulate cortex, insula, inferior frontal gyrus, middle frontal gyrus, hippocampus, precentral gyrus, and paracentral lobule; (c) and (d) represent mean GM maps for component 9 and 10, respectively. Connectogram shown in the middle for both FNC, left image for Figure 10a and right image for Figure 10c.

Figure 10c and connectogram of the right image illustrates positive connectivity in SC, CB, SM, along with a combination of some highly correlated positive and negative connections in cognitive control; GM also structurally increased in CC, SM, AU, and CB. Regions within the identified networks exhibit positive functional connectivity and share similar gray matter structural characteristics, this correlation suggests a relationship between functional dynamics and underlying structural architecture within the network. In addition, we observed that the CC exhibits greater interaction with other brain regions compared to other domains. We identified positive connectivity between insula, inferior frontal gyrus, middle frontal gyrus from CC and components precentral gyrus, paracentral lobule within SM. Besides, we observed decreased connectivity between components insula, inferior frontal gyrus, middle frontal gyrus, hippocampus from CC and posterior cingulate cortex from DM; between insula, inferior frontal gyrus, hippocampus of CC and component caudate of SC. The precentral gyrus of SM is negatively activated with middle frontal gyrus of CC and posterior cingulate gyrus of DM. The paracentral lobule is also negatively activated with the component caudate of SC.

## Discussion

The current identification of brain disorders heavily depends on observing clinical symptoms. While neuroimaging techniques offer a potentially more objective and biology-driven way to quantify brain abnormalities, the intricacies of the human brain and the presence of various sources of noise in neuroimaging signals pose challenges. In addition, multimodal brain imaging studies can provide a more complete understanding of the brain and its disorders (Calhoun & Sui, 2016). Numerous studies are currently underway, gathering multimodal brain imaging data and comprehensive information from the same participants. It is crucial to capitalize on these various imaging data types to extract complementary information. For instance, fMRI captures the dynamic hemodynamic response associated with neural activity in the brain, while sMRI allows us to estimate tissue types for each voxel in the brain, including GM, white matter and cerebrospinal fluid. Additionally, diffusion MRI offers insights into the integrity of white matter tracts and structural connectivity. The primary motivation behind concurrently analyzing multimodal data lies in harnessing the cross-information present in the dataset. Multimodal approaches unveil essential relationships that might remain undetected when relying on a single modality alone.

Multimodal methods are mostly limited to highly reduced summaries, e.g., a single map for fMRI like ALFF. Different imaging modalities may have varying spatial resolutions and anatomical landmarks. A single ALFF map may lack the detailed anatomical information needed for accurate alignment with other modalities, posing challenges in the fusion process. We are focused on incorporating information about multiple brain networks linked to brain structure. Our first work represents a step forward, but only focused on a limited number of networks including posterior DM, cerebellum, and SC. In this work, we present a significant extension by expanding our focus to include all 53 ICNs that allow us to evaluate network specific functions as well as inter-network connectivity information (functional network connectivity). This work presents a technique to fuse gray matter and multiple rest fMRI networks using a novel approach that combines concepts in group ICA and joint ICA. This approach minimizes those fusion challenges by combining the strengths of group ICA and parallel multilink jICA (Khalilullah et al., 2023). The inclusion of group ICA also facilitates the use of back-reconstruction to estimate spatial maps for individual brain networks, which was not part of the previous algorithm. These maps are derived from a functional-structural fusion algorithm, allowing for individual testing to assess group differences or associations with variables of interest in the Alzheimer disease data. In addition, we can compute the relationship among brain networks via FNC from the results. This provides an extremely rich set of multimodal and unimodal output that has not been available in prior approaches.

In this approach, we observed structural and functional alterations between healthy controls (HCs) and Alzheimer’s disease (AD) patients. Particularly noteworthy patterns in both structural and functional aspects were identified, with a focus on the auditory, visual, subcortical, cognitive control, and cerebellum regions that are recognized as being notably impacted areas for psychiatric disease, particularly for the Alzheimer’s disease (Agosta et al., 2023; Azeem A, 2023; Cai et al., 2015; Cheng et al., 2023; Elvira-Hurtado et al., 2023; Kawabata et al., 2022; Khalilullah et al., 2023; Kim et al., 2023; Lesh et al., 2011; Lin et al., 2017; Liu et al., 2018; Mavroudis, 2019; McEvoy et al., 2023; Sendi et al., 2023; Tang et al., 2021; Tarawneh et al., 2022; Tentolouris-Piperas et al., 2017; Xiao et al., 2022).

Based on our results, we find the thalamus, caudate, and putamen of the subcortical regions of the brain represent functionally lower connectivity in AD group as well as structurally reduced in AD. The subcortical regions act as pivotal sensory gates, facilitating the exchange of information with the cortical regions. The previous studies (Cai et al., 2015; Khalilullah et al., 2023; Kim et al., 2023; Lesh et al., 2011; Tentolouris-Piperas et al., 2017; Xiao et al., 2022) also found abnormalities in the psychiatric diseases, particularly for the schizophrenia and Alzheimer’s patients. Our analysis of FNC and mean GM spatial maps (Figure 9c-d) also suggests positive engagement of the caudate nucleus and thalamus with multiple domains. It appears that the basal ganglia (BG) network may be more directly linked to each of these regions. This finding represents a novel discovery not observed in our previous work. Additionally, we identified structural and functional reductions in cognitive control regions. Specifically, we observed structural reductions in the insula and inferior frontal gyrus in Alzheimer’s disease (AD), while the parahippocampal region, crucial for memory, showed both lower functional activity and structural reductions in AD. Emerging research indicates that the insula, inferior frontal gyrus of the cognitive control network stand as a pivotal hub within human brain networks and is identified as the most susceptible region to the effects of Alzheimer’s disease (AD) (Agosta et al., 2023; Lin et al., 2017; Liu et al., 2018; Schwab et al., 2020; Sendi et al., 2023), which are also in line with our research findings. In addition to the cognitive control regions of the brain, previous studies have shown that individuals with Alzheimer’s disease may experience difficulties in processing auditory information (Azeem A, 2023; Khalilullah et al., 2023; McEvoy et al., 2023; Tarawneh et al., 2022).

Structural and functional changes in the auditory domain were observed in our study. The superior temporal gyrus is structurally reduced whereas the middle temporal gyrus and inferior temporal gyrus are functionally increased in AD; the middle temporal gyrus of the auditory is also structurally increased in AD. These changes relative to controls may contribute to the deficits in auditory processing for the AD patients. In the visual network, the lingual gyrus and middle occipital gyrus show lower functional connectivity in AD. These results consistent with the previous studies (Elvira-Hurtado et al., 2023; Kawabata et al., 2022; Liu et al., 2017). We also notice structural-functional alternation in lingual gyrus and middle occipital gyrus of visual network. The changes observed in the structure and function of the visual network in Alzheimer’s disease can have a profound impact on information processing within the brain. This, in turn, may contribute to cognitive deficits and give rise to various clinical symptoms associated with Alzheimer’s disease.

In cerebellum regions, we find both increases and decreases functionally as well as structurally. Changes in the cerebellum may thus contribute to the cognitive symptoms observed in Alzheimer’s disease. The cerebellum is interconnected with various brain regions through extensive neural networks. Alterations in the cerebellum may impact these functional networks and contribute to the overall cognitive decline seen in Alzheimer’s disease. Emerging evidence suggests that the cerebellum, traditionally associated with motor function, may undergo structural and functional changes in Alzheimer’s disease (Cheng et al., 2023; Mavroudis, 2019; Tang et al., 2021). Understanding the extent and implications of cerebellar involvement is crucial for gaining a comprehensive understanding of the disease and may open new avenues for therapeutic interventions. Another interesting finding of our joint analysis is to separate overlapping regions of the brain networks which show opposite correlation from one another. Our findings include detailed spatial maps of altered activation patterns in key regions of the brain networks. This provides localized insights into the impact of AD on specific brain regions.

## Conclusions

In conclusion, our study presents a robust framework, pmg-jICA, for the joint analysis of multimodal MRI data, enabling the fusion of structural and functional information. By overcoming challenges in integrating different imaging modalities, our approach enhances the depth of understanding of brain alterations in Alzheimer’s disease, by linking together and jointly decomposing data while still preserving the richness of the information available. The investigation into structural and functional alterations between healthy controls and Alzheimer’s patients highlights notable patterns in regions such as the auditory, visual, subcortical, cognitive control, and cerebellum. By conducting a joint analysis with group ICA, it facilitates the comprehensive utilization of the variability among structural connections to specific functional networks and we can leverage the combined distribution of multimodal imaging data, ultimately enhancing our capability to discern between health and disease. The identified changes in connectivity and structure provide insights into the impact of Alzheimer’s disease on essential brain functions and behaviors. Moreover, the findings emphasize the importance of multimodal approaches in unraveling complex relationships that may remain elusive in unimodal studies. The study’s contributions extend beyond Alzheimer’s disease, offering a methodological advancement for the joint analysis of diverse neuroimaging data to enhance our understanding of brain disorders. Considering the current approach and the state of multimodal neuroimaging research, possible future directions may include: a) using temporal information of fMRI data for a more comprehensive understanding of brain dynamics; b) integrating additional modalities for better understanding of the underlying biological mechanisms associated with brain structure and function; c) incorporating improved machine learning and deep learning approaches to further optimize the integration of multimodal data.

## ACKNOWLEDGMENTS

This study was supported by National Science Foundation (NSF) 2316421, 2112455 and National Institutes of Health (NIH) RF1AG063153.

## DATA AVAILABILITY STATEMENT

The data are used from the Open Access Series of Imaging Studies (OASIS-3) and can be access via the following website https://www.oasis-brains.org.

