## Supplementary Material_1_2 for "Parallel Multilink Group Joint ICA: Fusion of 3D Structural and 4D Functional Data Across Multiple Resting fMRI Networks"

This table summarizes structure-function coupling.

Table II: Structure-function correlation

| | Modality | Brain Networks | Inference |
|---|---|---|---|
| **Component 2 of Combination-1** | **GM** | **AU:** Middle temporal gyrus | Increased in AD |
| | **fMRI** | Areas of CC | Increased in AD |
| | | **SC:** Caudate; Areas of CC | Lower in AD |
| **Component 4 of Combination-2** | **GM** | **AU:** Middle temporal gyrus | Increased in AD |
| | | **CC:** Insula, Inferior frontal gyrus | Lower inAD |
| | **fMRI** | **CC:** Nearly at hippocampal regions | Increased in AD |
| | | **SC:** Thalamus; VS: Lingual gyrus | Lower in AD |
| **Component 6 of Combination-3** | **GM** | **AU:** Superior temporal gyrus;<br>**SC:** Thalamus, Caudate;<br>**CC:** Insula, Inferior frontal gyrus | Lower in AD |
| | **fMRI** | **AU:** Superior temporal gyrus, Middle temporal gyrus | Increased in AD |
| | | **SC:** Caudate, Putamen;<br>**CC:** Parahippocampal | Lower in AD |
| **Component 6 of Combination-4** | **GM** | Areas of CC | Lower in AD |
| | **fMRI** | Caudate, Putamen | Lower in AD |
| **Component 8 of Combination-20** | **GM** | **SC:** Caudate, Thalamus;<br>Areas of CB | Lower in AD |
| | **fMRI** | Some areas of VS | Increased in AD |
| | | Some areas of VS | Lower in AD |
| **Component 8 of Combination-23** | **GM** | Areas of CB | Lower in AD |
| | **fMRI** | **VS:** Middle occipital gyrus<br>**AU:** Middle temporal gyrus | Lower in AD |
| **Component 6 of Combination-25** | **GM** | **AU:** Superior temporal gyrus;<br>**CC:** Insula, Inferior frontal gyrus, Parahippocampal gyrus | Lower in AD |
| | **fMRI** | **AU:** Middle temporal gyrus, Inferior temporal gyrus | Increased in AD |
| | | Areas of VS | Lower in AD |
| **Component 3 of Combination-28** | **GM** | **AU:** Superior temporal gyrus;<br>**SC:** Thalamus, Caudate;<br>**CB:** Declive, Uvula, Tuber | Lower in AD |
| | **fMRI** | **CC:** Superior frontal gyrus, Medial frontal gyrus | Increased in AD |
| | | **CC:** Middle frontal gyrus | Lower in AD |
| **Component 3 of Combination-42** | **GM** | Areas of CB | Increased in AD |
| | **fMRI** | **AU:** Middle temporal gyrus<br>**DMN:** Posterior Cingulate | Increased in AD |
| | | **CC:** Middle frontal gyrus, Medial frontal gyrus, Supramarginal gyrus; | Lower in AD |

| | | | |
|---|---|---|---|
| **Component 4 of Combination-51** | **GM** | Areas of CB | Lower in AD |
| | **fMRI** | Areas of VS; Areas of CB | Increased in AD |
| **Component 2 of Combination-53** | **GM** | **AU:** Inferior temporal gyrus; Areas of CB | Lower in AD |
| | **fMRI** | Areas of VS; Areas of CB | Increased in AD |
| | | Areas of CB | Lower in AD |
