## Supplementary Material_2_2 for "Parallel Multilink Group Joint ICA: Fusion of 3D Structural and 4D Functional Data Across Multiple Resting fMRI Networks"

In Figure 11a, FNC shows increased connectivity within SC, VS, CC, and CB. The SC and CC components are more activated than VS and CB. In inter-network connectivity, we observed heightened connectivity between hippocampus from CC and postcentral gyrus within SM; between anterior DM and superior parietal lobule from SM. GM shows increased connectivity in AU, VS, and CB. In Figure 11c, the more activated regions in SC and VS within the networks. We also identify inter-network connectivity between hippocampus from CC and postcentral gyrus within SM; between inferior frontal gyrus from CC and putamen within SC; between middle occipital gyrus from VS and postcentral gyrus within SM. GM exhibits positive pattern in VS, AU, and CB.

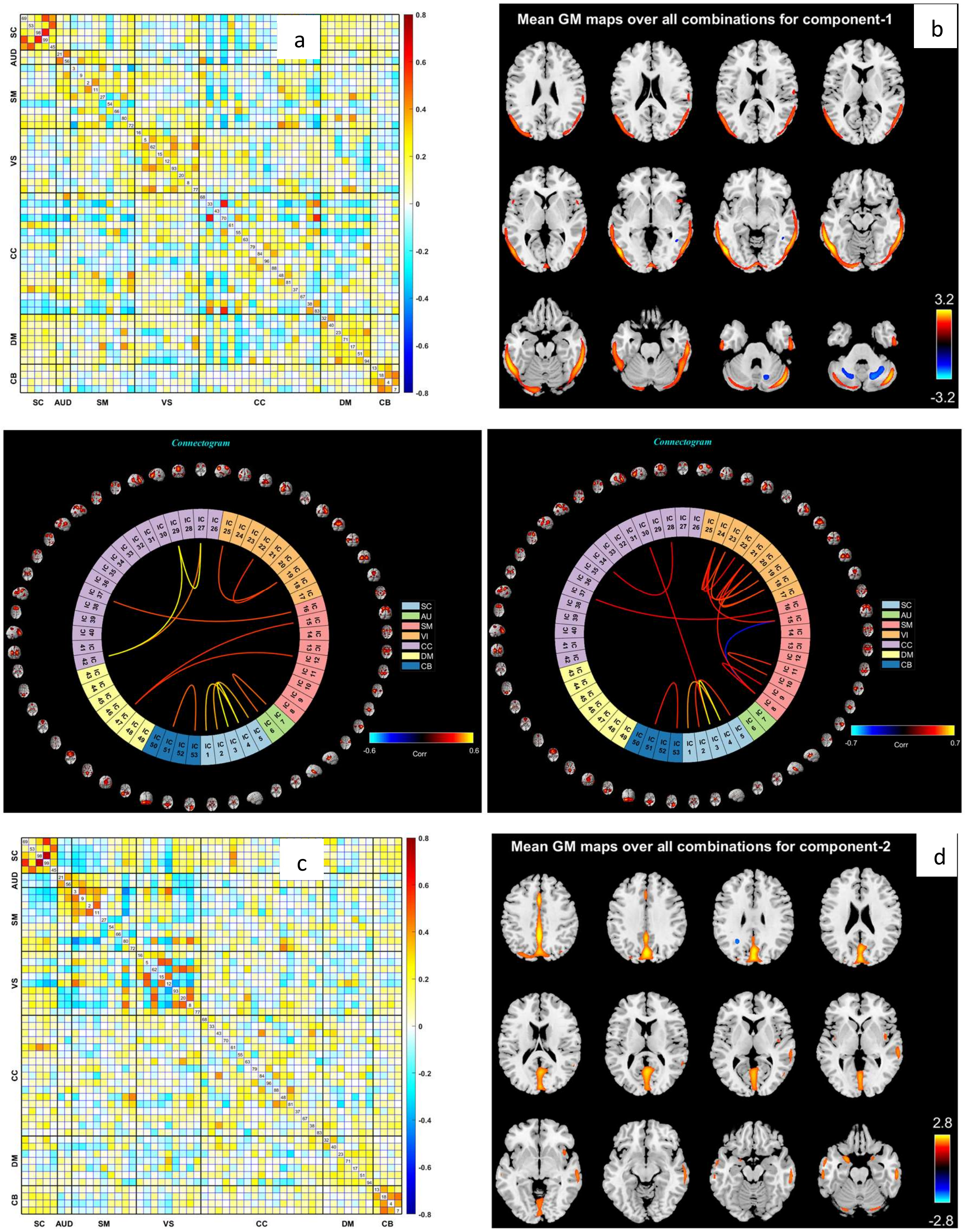


Fig. 11: FNC from reconstructed component 1 and 2 of all ICNs; highly correlated regions are (a) hypothalamus, thalamus, caudate, middle cingulate cortex, middle temporal gyrus, inferior parietal lobule, insula, inferior frontal gyrus, anterior cingulate cortex, and superior parietal lobule, (c) hypothalamus, thalamus, caudate, middle cingulate cortex, hippocampus, left inferior parietal lobule, superior parietal lobule, postcentral gyrus, lingual gyrus, and middle temporal gyrus; (b) and (d) show mean GM maps for component 1 and 2, respectively.

In Figure 12a, we show highly positive connectivity regions in SC and SM, while SC shows a mixture of connections. In inter-network connectivity, there are increased connections between superior parietal lobule from AM, middle occipital gyrus within VS and anterior DM; between inferior parietal lobule, superior medial frontal gyrus from CC and middle temporal gyrus; between superior medial frontal gyrus, supplementary motor area from CC and anterior DM. In Figure 12c, we found increased functional connectivity within SC and VS, they are more activated than other regions such as SM, CB, DM, and CC. In addition to intra-network connectivity, we identify increased connectivity between inferior frontal gyrus of CC and precuneus within DM; between hippocampus of CC and postcentral gyrus from SM; between precuneus of DM and hypothalamus of SC.

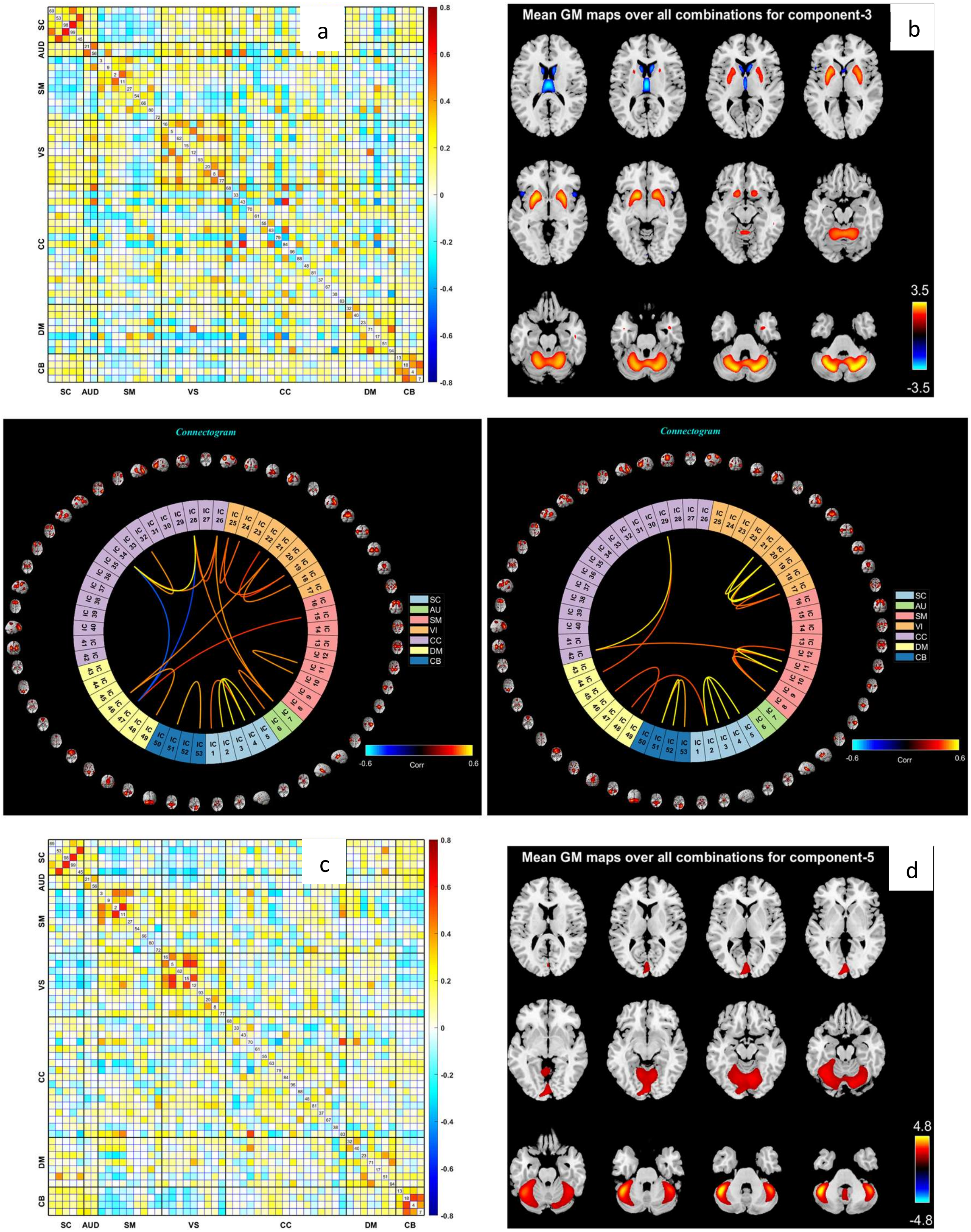


Fig. 12: FNC from reconstructed component 3 and 5 of all ICNs; highly correlated regions are (a) inferior parietal lobule, insula, superior frontal gyrus, superior medial frontal gyrus, supplementary motor area, middle temporal gyrus, postcentral gyrus, superior parietal lobule, middle occipital gyrus fusiform gyrus, posterior cingulate cortex, anterior cingulate cortex, and SC domain, (c) precuneus, posterior cingulate cortex, inferior frontal gyrus, hippocampus, postcentral gyrus, paracentral lobule, middle occipital gyrus, middle temporal gyrus, cuneus, thalamus, hypothalamus, putamen; (b) and (d) represent mean GM maps for component 3 and 5, respectively.

In Figure 13a, we identify increased intra-network connectivity in SC, VS, SM, CB, CC, and DM. The SC and VS are highly activated than others, while there is a mixture connection within VS. In inter-network connectivity, we found increased connectivity between inferior frontal gyrus of CC and precuneus from DM; between inferior parietal lobule, inferior frontal gyrus from CC and inferior occipital gyrus within VS; between posterior DM and inferior occipital gyrus from VS. In Figure 13c, we observed increased intra-network connectivity within SC, VS, also in SM and CC. We also observed inter-network connectivity between superior parietal lobule within SM, fusiform gyrus in VS and anterior DM; between Hippocampus from CC and right postcentral gyrus within SM; between left inferior parietal lobule from CC and postcentral gyrus within SM.

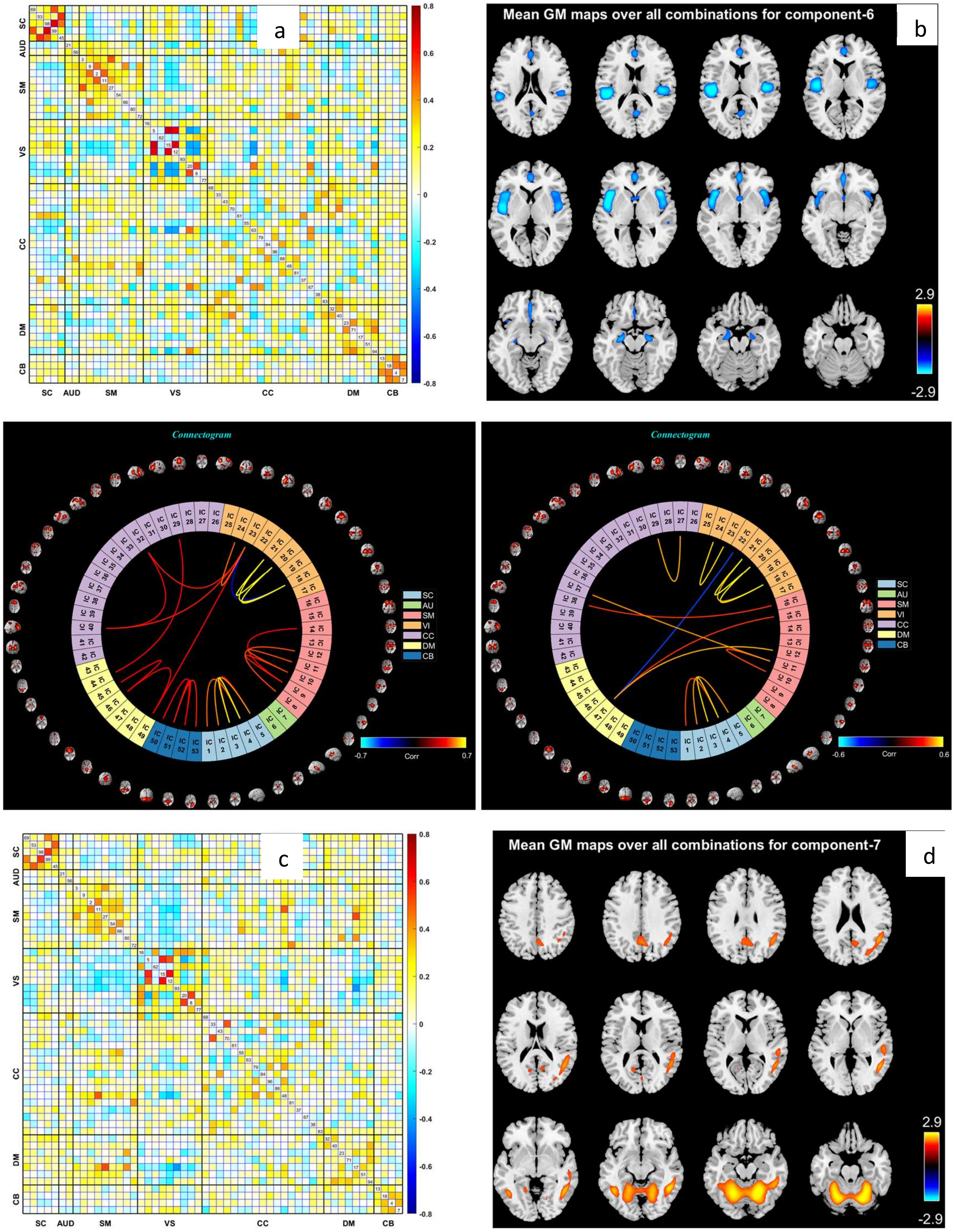


Fig. 13: FNC from reconstructed component 6 and 7 of all ICNs; highly correlated regions are (a) hippocampus, left inferior parietal lobule, inferior frontal gyrus, posterior cingulate cortex, postcentral gyrus, inferior occipital gyrus, lingual gyrus, (c) hippocampus, postcentral gyrus, superior parietal lobule, anterior cingulate cortex, cuneus, occipital gyrus, fusiform gyrus; (b) and (d) represent mean GM maps for component 6 and 7, respectively.
